# Multi-protein silencing using WRAP-based nanoparticles: a proof of concept

**DOI:** 10.1101/2025.02.07.637024

**Authors:** Karidia Konate, Irène Pezzati, Karima Redjatti, Estelle Agnel, Eric Vivès, Sandrine Faure, Pascal de Santa Barbara, Prisca Boisguérin, Sébastien Deshayes

## Abstract

Cancer remains the leading cause of death, with chemotherapy, radiotherapy, and surgical resection being the primary treatment methods. However, chemotherapy’s side effects, surgical limitations, and drug resistance present significant challenges. Small interfering RNA (siRNA) has emerged as a promising tool in cancer therapy due to its ability to silence disease-related genes selectively. Recent advancements in non-viral delivery systems, particularly cell-penetrating peptides (CPPs), have enhanced the efficacy of siRNA delivery. The use of siRNA as a therapeutic tool in cancer treatment has been reported. However, silencing only one target protein has only minor effects on tumor cell proliferation as previously shown for WRAP-based nanoparticles targeting cyclin-dependent kinase 4 (CDK4) in human glioblastoma cells. Here, we designed a more sophisticated approach to enhance therapeutic efficacy, encapsulating multiple siRNAs targeting CDK4, cyclin D1 (CD1), and Mcl-1 proteins. The siRNA cocktail, delivered via WRAP5 nanoparticles, effectively silenced these targets and reduced cell proliferation in human glioblastoma cells. Furthermore, the nanoparticles also demonstrated potential therapeutic impact in gastrointestinal stromal tumors (GIST), a rare cancer characterized by its tendency to resist standard treatments. This study highlights the versatility of WRAP5 nanoparticles as a platform for personalized cancer therapy, suggesting that siRNA delivery systems may be tailored to specific cancer types for more effective treatment strategies.

## INTRODUCTION

Based on the global cancer statistics database provided by the Global Cancer Observatory (GLOBOCAN) estimation for 2022, cancer is the first leading cause of death worldwide, accounting for almost 10 million deaths per year (nearly one in six deaths).^1^ Despite the considerable progress made in the field of anti-cancer therapy, particularly in the development of chemotherapy, radio-chemotherapy, and hormonal therapy, surgical resection in combination with chemo-radiotherapy remains the established method for treating malignant cancers.^2^ However, chemotherapy is not without its drawbacks, as evidenced by the occurrence of adverse side effects and the potential for inadvertent damage to healthy cells. Moreover, the inadequacy of surgical resection can result in cancer recurrence in a variety of cases, including glioblastoma multiform (GBM).^3^ In addition, the primary challenge to the effective treatment of cancer patients concerns the appearance of drug resistance as observed for gastrointestinal stromal tumors (GIST) mainly through secondary mutations.^4^

Consequently, novel therapeutic approaches are now being developed to target the fundamental mechanisms and acquired capacities that transform healthy cells and tissues into cancerous tumors. In this context, a particularly fruitful area of research in the domain of precision medicine employs small interfering RNA (siRNA).^2,5,6^ In detail, the potential efficacy of siRNA in treating diseases is attributable to the binding to specific mRNA sequences resulting in an efficient silencing of disease-related genes. This was further demonstrated by the fact that six siRNAs have been approved by the FDA (US Food and Drug Administration) and the EMA (European Medicines Agency).^7^ All six of these siRNAs target mRNAs made in liver cells, with five of them delivered as GalNAc conjugates and one as siRNA formulated in lipid-based nanoparticles (LNP)^8,9^, named Patisiran.^10^

To target other organs or cancer tissues, a range of other delivery systems have been designed in different laboratories. These include non-viral carriers composed of polymers, lipids, or inorganic materials, which are to be tested in preclinical and clinical studies.^11^ However, for some of them, several side effects were observed such as the activation of Fas Ligand-mediated cell death by polyethylenimine (PEI) or toxicity and platelets and CD11-b-positive cells aggregation in the lung by poly-(L-lysine) (PLL).^12^

In the field of safer delivery systems, cell-penetrating peptides (CPPs, comprising up to 30 amino acids) have been proposed as an up-and-coming alternative. Notably, those CPPs capable of forming nanoparticles in the presence of siRNAs are particularly interesting.^13–15^ This non-covalent strategy is based on electrostatic and hydrophobic interactions between both the peptide and the siRNA, thus opening peptides to the field of nanomedicine.

In this context, we designed a family of amphipathic peptides called WRAP (W- and R-rich Amphipathic Peptides) able to form peptide-based nanoparticles (PBNs) once incubated with siRNA at a specific CPP:siRNA molar ratio (MR).^16^ The high efficacy of the WRAP-based PBNs in delivering siRNAs across different cancer cell lines can be attributed to their rapid cellular internalization (~15 minutes) mainly through direct membrane translocation, with only a minor fraction entering via endocytosis-dependent pathways.^17^ Based on a structure-activity relationship (SAR) study, we identified two lead peptides WRAP1 (LLWRLWRLLWRLWRLL) and WRAP5 (LLRLLRWWWRLLRLL) showing nearly the same delivery properties.^18^ Compared to lipid-based reagents for siRNA transfection, we demonstrated that WRAP:siRNA nanoparticles are as efficient for protein silencing without cytotoxicity.^18^ Finally, the first *in vivo* investigation performed with WRAP:siRNA nanoparticles demonstrated effective luciferase silencing in a mouse xenograft model using glioblastoma cancer cells that overexpress luciferase.^19^

In our previous works, we focused mainly on glioblastoma (GBM), which represents the most prevalent form of malignant brain and other central nervous system (CNS) tumors. The incidence of glioblastoma is 3.21 per 100,000 population.^20^ Characteristics of this cancer were dysregulation of signal pathways and an aberrant cell cycle due to cyclin-dependent kinase 4 (CDK4) amplification or overexpression as reported for most of cyclin-dependent kinases (CDKs) in human cancers.^21,22^ Therefore, we aimed to target GBM cell proliferation by WRAP:siRNA nanoparticles downregulating CDK4 to develop a pharmacological treatment. However, despite observing significant CDK4 protein silencing, no phenotypical changes, such as a reduction in proliferation, were induced.^16,18^ One potential explanation for this phenomenon is that the inhibition of one single protein is not sufficient to arrest the whole cell cycle.

The objective of this study was to develop a more sophisticated approach to simultaneously target multiple proteins through the encapsulation of multiple siRNAs in the WRAP5-based nanoparticles. Specifically, we aimed to target both the cell cycle and cell survival, to inhibit cell proliferation and direct cancer cells to death. To target the cell cycle, we selected CDK4 as previously described ^16,18^ and its activator, the cyclin D1 (CD1) whose overexpression resulted in dysregulated CDK activity, rapid cell growth, bypass of key cellular checkpoints and ultimately cancer progression.^23^ To target cell survival, we selected the myeloid leukemia cell differentiation protein MCL-1, a key anti-apoptotic protein within the BCL-2 family.^24^ Indeed, it has been established that many cancers evaded pro-apoptotic stress signals by upregulating anti-apoptotic proteins, such as MCL-1, to maintain their survival.^25^ Furthermore, MCL-1 overexpression is often linked to therapeutic resistance.

In the present study, we reported on the encapsulation of three siRNAs targeting CDK4, CD1, and MCL-1 proteins as a potential cancer treatment. Firstly, we confirmed that the three proteins could be silenced individually by the WRAP5:siRNA nanoparticles in human glioblastoma U87 cells. Subsequently, we performed knockdown using different ratios of 2 or 3 encapsulated siRNAs. Following this screening, we demonstrated that WRAP5 nanoparticles encapsulating the three siRNAs with a defined ratio of 14/3/3 for siCD1/siCDK4/siMCL-1 (=siCOCK) silenced the three proteins by more than 80% in U87 cells. Furthermore, this particular siRNA cocktail was found to have a physiological impact on cell proliferation, in comparison to the nanoparticles encapsulating a single siRNA.

Furthermore, to evaluate if the WRAP5:siCOCK nanoparticles could be applied to other cancers, we analyzed them in gastrointestinal stromal tumors (GIST) which are rare cancers but represent the most common form of sarcoma of the digestive tract, with an incidence of 0.82 per 100,000 population.^26^ GISTs develop predominantly in the stomach and are caused in approximately 75% of cases by gain-of-function mutations in the gene of the receptor tyrosine kinase c-KIT. Imatinib (IM), a tyrosine kinase inhibitor (TKI), is the standard first-line treatment for GIST. However, it is important to note that complete remission is rare, and resistance is frequent, highlighting the need for the development of novel therapeutics.

As was the case for the glioblastoma U87 cells, we first analyzed the silencing of the three proteins individually and then all together using WRAP nanoparticles encapsulating single siRNA or the siCOCK. Interestingly, we found a similar effect on cell proliferation when the WRAP5:siCOCK was applied on GIST cells, however with another siRNA ratio. These results suggest that the efficacy of cancer treatment using siRNAs should be adapted according to the specific type of cancer being treated. This approach could represent a significant advancement in the field of personalized cancer therapy.

## MATERIALS and METHODS

### Materials

WRAP5 peptide was synthesized at the SynBio3 platform (IBMM Montpellier) and the crude product was purified in house following a qualitative analysis by HPLC/MS (~95% purity). The different siRNA sequences were purchased from Eurogentec (See **Table S1**). The siRNA stock solutions were prepared in RNase-free water.

### WRAP5:siRNA nanoparticle formulation

WRAP5 solutions were prepared in pure water (Sigma-Aldrich). Nanoparticles were formulated in pure water supplemented by 5% glucose (Sigma-Aldrich) by mixing equal volumes of siRNA and WRAP5 at the corresponding molar ratio (R = 20) at room temperature. The molar ratio R = 20 corresponded to a 20-fold higher molar concentration of WRAP5 compared to the siRNA as already described.^16^ For WRAP5:siCOCK nanoparticles, siRNA cocktails were pre-mixed according to siRNA stoichiometry: 80% siCD1 and 20% siCDK4 for 4/1 “binary cocktail” and 70% siCD1, 15% siCDK4 and 15% siMCL-1 for 14/3/3 “tertiary cocktail” in human U87 glioblastoma cells as well as 30% siCD1, 40% siCDK4 and 30% siMCL-1 for 6/8/6 “tertiary cocktail” in human GIST-T1 cells.

### Dynamic light scattering (DLS)

WRAP5:siRNA nanoparticles (WRAP5 = 10 μM, siRNA = 500 nM, R = 20) were evaluated with a Zetasizer NanoZS (Malvern) in terms of mean size (Z-average) of the particle distribution and of homogeneity (PdI). All results were obtained from three independent measurements (three runs for each measurement at 25°C).

### Culture conditions

Human glioblastoma cells (U87) were grown in a complete medium: DMEM with GlutaMAX™ (Thermo Fisher Scientific), penicillin/streptomycin (Thermo Fisher Scientific), 10% fetal bovine serum (FBS, Thermo Fisher Scientific), non-essential amino acids NEAA 1 X (Thermo Fisher Scientific) and 0.1% hygromycin (Invitrogen). Human gastrointestinal stromal tumor (GIST-T1) cells were grown in a complete medium: DMEM with GlutaMAX™ (Thermo Fisher Scientific), 1% penicillin/streptomycin (Thermo Fisher Scientific), 10% fetal bovine serum (FBS, Thermo Fisher Scientific). All the cells were maintained in a humidified incubator with 5% CO2 at 37°C.

### Transfection experiments

For Western blot assays, 75,000 U87 and 150,000 GIST-T1 cells were seeded 24 h before experiment into 24-well plates. For nanoparticle incubation, the cells were incubated with 175 μL of fresh pre-warmed serum-free DMEM + 75 μL of the nanoparticle solutions at the indicated concentrations or 75 μL 5% glucose for the non-treated cells. After 1.5 h of incubation, 250 μL DMEM supplemented with 20 % FBS (final FBS concentration = 10 %) were added to each well without withdrawing the transfection reagents. Cells were then incubated for another 24 h and finally lysed for Western blot evaluation.

For clonogenic assays, 350 U87 or GIST-T1 cells were seeded 24 h before the experiment into 6-well plates. For nanoparticle incubation, the cells were incubated with 1.7 mL of fresh pre-warmed serum-free DMEM + 300 μL of the nanoparticle solutions at the indicated concentrations or 300 μL 5% glucose for the non-treated cells. After 1.5 h of incubation, the medium was removed and replaced by 2 mL DMEM supplemented with 10 % FBS. Cells were then incubated for 14 days.

For confocal microscopy, 300,000 GIST cells were seeded 24 h before the experiment into a glass bottom dish (FluoroDish, 35 mm). For nanoparticle incubation, the cells were incubated with 1800 μL of pre-warmed serum-free DMEM FluoroBrite + 200 μL of the nanoparticle solutions at the indicated concentrations or 200 μL 5% glucose for the non-treated cells.

### Cell cytotoxicity measurement

The cytotoxicity induced by the nanoparticles was evaluated using Cytotoxicity Detection Kit^Plus^ (LDH, Sigma-Aldrich) following the manufacturer’s instructions. At least one well of the 24-well plate was used as LDH positive control (100% toxicity) by adding Triton X-100 (Sigma-Aldrich) to a final concentration of 0.1% (~15 min incubation at 37°C). Afterward, 50 μL supernatant of each well were transferred to a new clear 96-well plate. 50 μL of the “dye solution/catalyst” mixture was added to the supernatant and incubated in the darkness for 30 min at room temperature. The reaction was stopped by adding 25 μL of HCl (1 N) to each well before measuring the absorption at 490 nm. Relative toxicity (%) = [(exp. value – value non-treated cells) / (value triton – value non-treated cells)] x 100.

### Western blotting

Transfected cells were washed in PBS, and lysed in RIPA buffer [50 mM Tris pH 8.0, 150 mM sodium chloride, 1% Triton X-100, 0.1% SDS (sodium dodecyl sulfate, Sigma-Aldrich), including protease inhibitors (SigmaFAST, Sigma-Aldrich)]. Cells were incubated for 5 min on ice with 130 μL/24-well lysis buffer. Thereafter, cells were scraped and transferred in a 1.5 mL tube. After 5 min on ice, the cell lysates were centrifuged (10 min, 16,100 g, 4°C), supernatants were collected and protein concentrations were determined using the Pierce BCA Protein Assay (Thermo Fisher Scientific). Cell extracts were separated by 4-20% Mini-PROTEAN® TGXTM Precast Gel (Bio-Rad). After electrophoresis, samples were transferred onto Trans-Blot® Turbo™ Mini PVDF Transfer membrane (Bio-Rad). As antibodies, we used anti-CDK4 rabbit mAb D9G3E, anti-MCL-1 rabbit, anti-Vinculin rabbit mAb E1E9V, anti-mouse IgG HRP and anti-rabbit IgG HRP(all from Cell Signaling) as well as anti-CD1 (SP4) (Thermo Fisher Scientific). Blots were revealed with the Pierce ECL plus Western blotting substrate (Thermo Fisher Scientific) on an Amersham imager 600 (GE Healthcare Life Science) or a Sapphire RGB & NIR Biomolecular Scanner (Azure biosystems). The signal intensities of the blots were quantified using Fidji ImageJ software. Each band intensity corresponding to a distinguished condition is then normalized to the band intensity of non-treated cells (N.T.) (= 100%): Relative Signal Intensity (%) = intensity (condition x)/intensity (N.T.) x 100.

### Clonogenic assay

The medium of the transfected U87 cells was removed and 1.5 mL/well of methanol:acetic acid solution (3/1, vol/vol) was added to fix the cell colonies. After 20 min incubation, the solution was replaced by 1.5 mL/well of Giemsa (Sigma-Aldrich)/H_2_O (3.5/10, vol/vol) for 20 min. The medium of the transfected GIST-T1 cells was removed and cells were washed two times with PBS1X. Then cells were fixed using 0.5 mL/well of 4% paraformaldehyde (PFA) solution. After 10 min incubation, cells were rinsed two times with PBS1X, and 0.5 mL/well of crystal violet solution (in 10% Ethanol) was added to color cell colonies for 10 min. Finally, the dye solution was removed and the wells were rinsed many times with H_2_O or PBS1X until the background of the wells was clear. After air-drying the wells, an image of the whole 6-well plate was acquired on an Amersham imager 600 (GE Healthcare Life Science) or a Sapphire RGB & NIR Biomolecular Scanner (Azure biosystems). Proliferation (%) = number of colonies (condition x)/number of colonies (N.T.) x 100.

### Confocal microscopy

Transfected cells were incubated for 10 min Hoechst 33342 dye (Sigma-Aldrich) for nucleus labeling. Afterward, cells were washed twice with D-PBS and covered with FluoroBrite DMEM medium (Thermo Fisher Scientific). Live cell images were acquired on an inverted Zeiss LSM800 microscope using a lens Apo 63x/1.2 W DICIII. All confocal acquisitions were performed using the following diode lasers: 405 nm with a Hoechst filter (400 nm - 456 nm), 488 nm with an Alexa488 filter (500 nm - 542 nm), 561 nm with an Atto550 filter (560 nm - 617 nm) and 640 nm with an Atto655 filter (656 nm - 700 nm). Image acquisition was done sequentially to minimize crosstalk between the fluorophores. Each confocal image was merged and adjusted with the same brightness and contrast parameters using the Fidji ImageJ software.

## RESULTS

### Targeting three different proteins using WRAP5:siRNA in U87 cells

As previously revealed by confocal microscopy, WRAP5-based nanoparticles are a suitable delivery system to transfect siRNAs in human U87 glioblastoma cells.^16,17^ However, our studies using WRAP5-loaded with siRNA targeting CDK4 did not reveal the expected phenotypic activity in terms of cancer cell proliferation reduction.^16,18^ We assumed that reducing the expression of only one targeted protein involved in the cell cycle regulation was not sufficient to disturb cell growth because other cyclin-dependent kinases (e.g. CDK2, CDK6) could compensate the CDK4 activity.^27,28^ To overcome this fact, we decided to silence simultaneously different proteins of the cell cycle such as CDK4 and Cyclin D1 (CD1) as well as the anti-apoptotic protein MCL-1 to stop the cell cycle progression and reduce cell survival.

First, we evaluated the dose-dependent silencing capacity of three different siRNAs especially designed to target proteins CD1, CDK4, and MCL-1 (siCD1, siCDK4, and siMCL-1) separately. For that, one of the three siRNAs was encapsulated by the WRAP5-based nanoparticle and applied to human U87 glioblastoma cells (**Figure 1A**). We used the same WRAP5:siRNA molar ratio between the peptide and the siRNA (molar ratio 20:1) which was described as optimal for siRNA delivery.^16,17,19^

**Figure 1:**
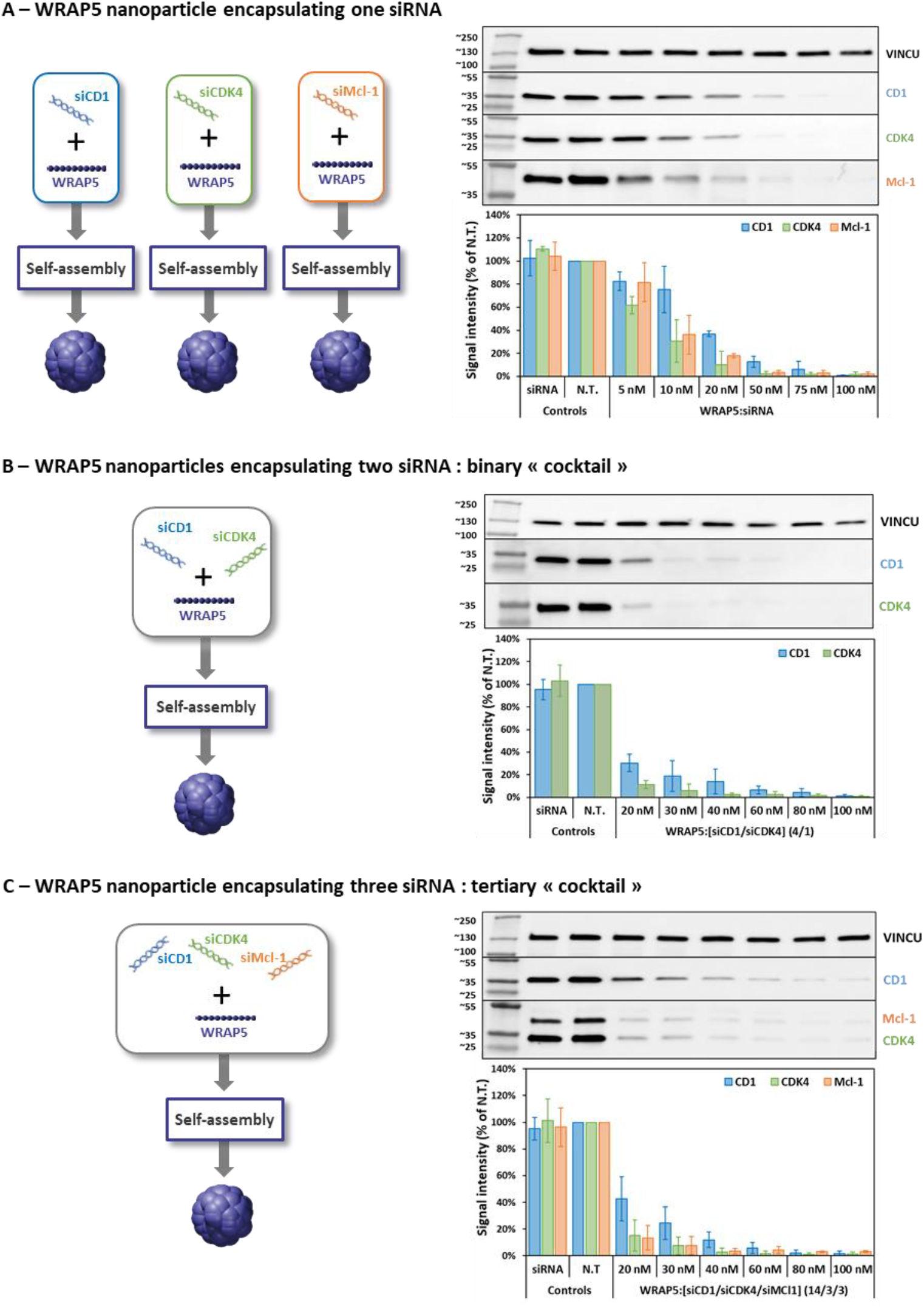
Evaluation of the dose-dependent silencing of WRAP5:siRNA nanoparticles in U87 cells. **(A)** WRAP5 nanoparticles encapsulating one siRNA (schematic representation on the left side) induced the dose-dependent silencing of CD1, CDK4, and MCL-1 proteins in human U87 glioblastoma cells at the indicated siRNA concentrations revealed by Western Blot after a 24 h incubation. **(B)** WRAP5 nanoparticles encapsulating two siRNAs (schematic representation on the left side) in a siRNA ratio 4/1 induced the dose-dependent simultaneous silencing of two proteins (CD1/CDK4) in human U87 glioblastoma cells at the indicated siRNA concentrations revealed by Western Blot after a 24 h incubation. **(C)** WRAP5 nanoparticles encapsulating three siRNAs (schematic representation on the left side) in a siRNA ratio 14/3/3 induced the dose-dependent simultaneous silencing of three proteins (CD1/CDK4/MCL-1) in human U87 glioblastoma cells at the indicated siRNA concentrations revealed by Western Blot after a 24 h incubation. Data represent mean ± SD, with n = 2 independent experiments in duplicates. Controls: Non-treated cells (N.T.) and siRNA alone.

WRAP5 nanoparticles encapsulating the siCDK4 revealed an important knock-down of the CDK4 protein expression (>80% using 20 nM siCDK4) as observed previously ^16^ as well as for the MCL-1 protein (~80% using 20 nM siMCL-1) (**Figure 1A**). Curiously, a higher siCD1 concentration (50 nM) was required to obtain the same knock-down efficiency (~80%). This was not explained by different sizes of the nanoparticles (all between 75-100 nm, see **Table 1**) but probably by an increased stability, a longer turn-over or a higher expression of the CD1 protein compared to CDK4 and MCL-1 proteins in U87 cells. CD1 was described as a key protein involved in different cancers ^29,30^, suggesting that its expression fluctuated according to cell types. This latter observation could explain why the CD1 silencing needed more than twice the siRNA concentration of the other targets (CDK4, MCL-1).

**Table 1:**
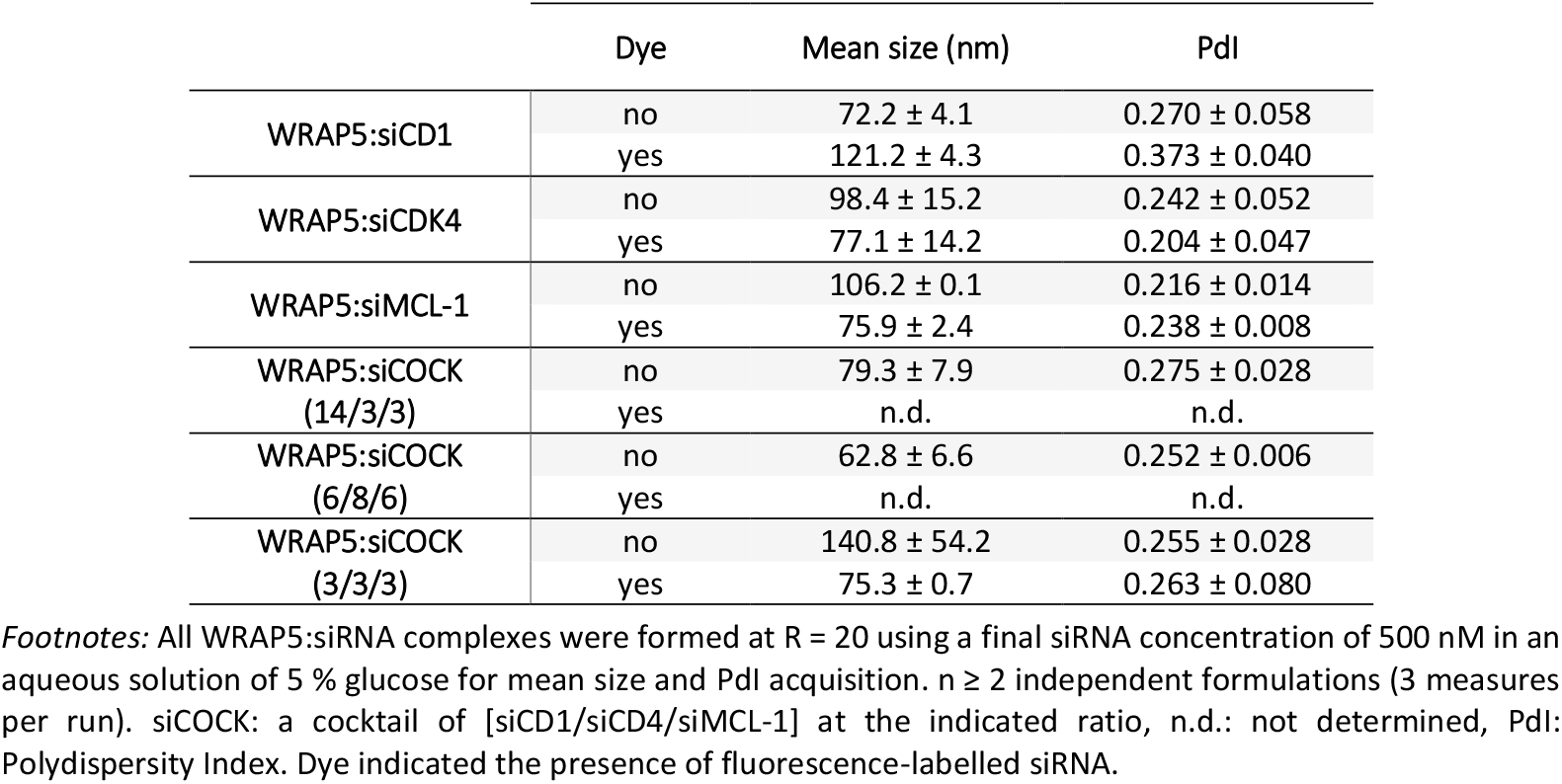
WRAP5 nanoparticles characterization by DLS measurements.

### WRAP5 nanoparticles encapsulating up to 3 siRNA for protein silencing in U87 cells

To avoid using high concentrations of CD1 siRNA, we decided to perform a “binary cocktail” approach by encapsulating together siCD1 and siCDK4 into one nanoparticle. With this condition, we expected the silencing of both proteins. To obtain a simultaneous and equivalent silencing of both proteins, we combined 80% of siCD1 with 20% of siCDK4, corresponding to a final ratio of 4/1 [siCD1/siCDK4] into WRAP5 nanoparticles (**Figure 1B**). As described below, the molar ratio between WRAP5 and siRNA was 20/1, and this will be kept constant for all following experiments independently if one, two, or three siRNAs were encapsulated.

By looking in detail at the “40 nM” condition corresponding to a nanoparticle encapsulating 32 nM siCD1 and 8 nM siCDK4, three important points could be revealed in human U87 glioblastoma cells. First, we showed an important dose-dependent simultaneous silencing of CD1 and CDK4 using this 4/1 siRNA ratio. Secondly, we observed a synergic effect as the knock-down as CDK4 silencing was higher (>90%) compared to the single siRNA-loaded nanoparticles at a similar concentration (~70% for WRAP5:siCDK4 at 10 nM) (**Figure 1A**). Interestingly, we also illustrated a slight synergic effect for the CD1 protein with a knock-down of ~80% in the binary condition (siRNAs at 30 nM with siCD1 at 24 nM and siCDK4 at 6 nM, **Figure 1B**) compared to the single-loaded nanoparticle (siCD1 at 50 nM, Fig.1A).

We observed similar behavior for the encapsulation of a “binary cocktail” targeting both CD1 and MCL-1 at a final siRNA ratio of 4/1 [siCD1/MCL-1] (**Figure S1**). This led us to extend the idea of siRNA combination, considering a “tertiary cocktail” formulation.

Knowing that a higher siCD1 concentration was needed to obtain the same silencing efficiency as CDK4 and MCL-1, we encapsulated the three different siRNAs in a “tertiary cocktail” using 70% siCD1, 15% siCDK4 and 15% siMCL-1, corresponding to a siRNA ratio of 14/3/3 [siCD1/siCDK4/siMCL-1] (**Figure 1C**) and we evaluated their effect in a dose-dependent manner in human glioblastoma U87 cells.

Then, the “20nM” condition was selected to assess the synergistic effects between the three knock-downs, which corresponded to 14nM siCD1, 3nM siCDK4, and 3nM siMCL-1. Interestingly, we observed a positive effect of the transfection of the three siRNAs because the effects of the “tertiary cocktail” were higher on the three proteins (70% for CD1, 90% for CDK4 and 80% for MCL-1) compared to the corresponding single siRNA-loaded nanoparticles at the same concentrations (~25% for WRAP5:siCD1 at 10 nM, 40% for WRAP5:CDK4 at 5 nM and 20% for WRAP5:MCL-1 at 5 nM – **Figure 1A**).

Finally, we evaluated the WRAP5 nanoparticle loaded with the siRNAs cocktail [siCD1/siCDK4/siMCL-1] (= WRAP5:siCOCK) compared to single siRNA-loaded nanoparticles by Dynamic Light Scattering (DLS, **Table 1**). The measurements revealed that WRAP5 encapsulating one or three siRNAs all formed nanoparticles with mean size between 70 and 100 nm with a PdI <0.3. Graphical representations of the measured mean sizes are visualized in **Figure S2**. Moreover, these nanoparticles were stable over 7 days when stored at 4°C (data not shown).

### WRAP5-based siRNA cocktail: optimal dosing for silencing in U87 cells

The next step was the evaluation of the WRAP5:siCOCK nanoparticles for their silencing efficiency compared to a negative control condition (WRAP5 encapsulating an inactive siRNA = siNEG). In the same experiment, we also compared the WRAP5:siCOCK with nanoparticles with one siRNA substituted by the siNEG to estimate if a synergic effect could be shown (**Figure 2A**). Two final siRNA concentrations (40 nM, 60 nM) were selected for this screening within the middle range of our previous dose response (**Figure 1C**) to ensure sufficient concentration for inhibiting target proteins. As reported before for PBNs, these concentrations were known to have no cytotoxic effects.^16,18,31,32^

**Figure 2:**
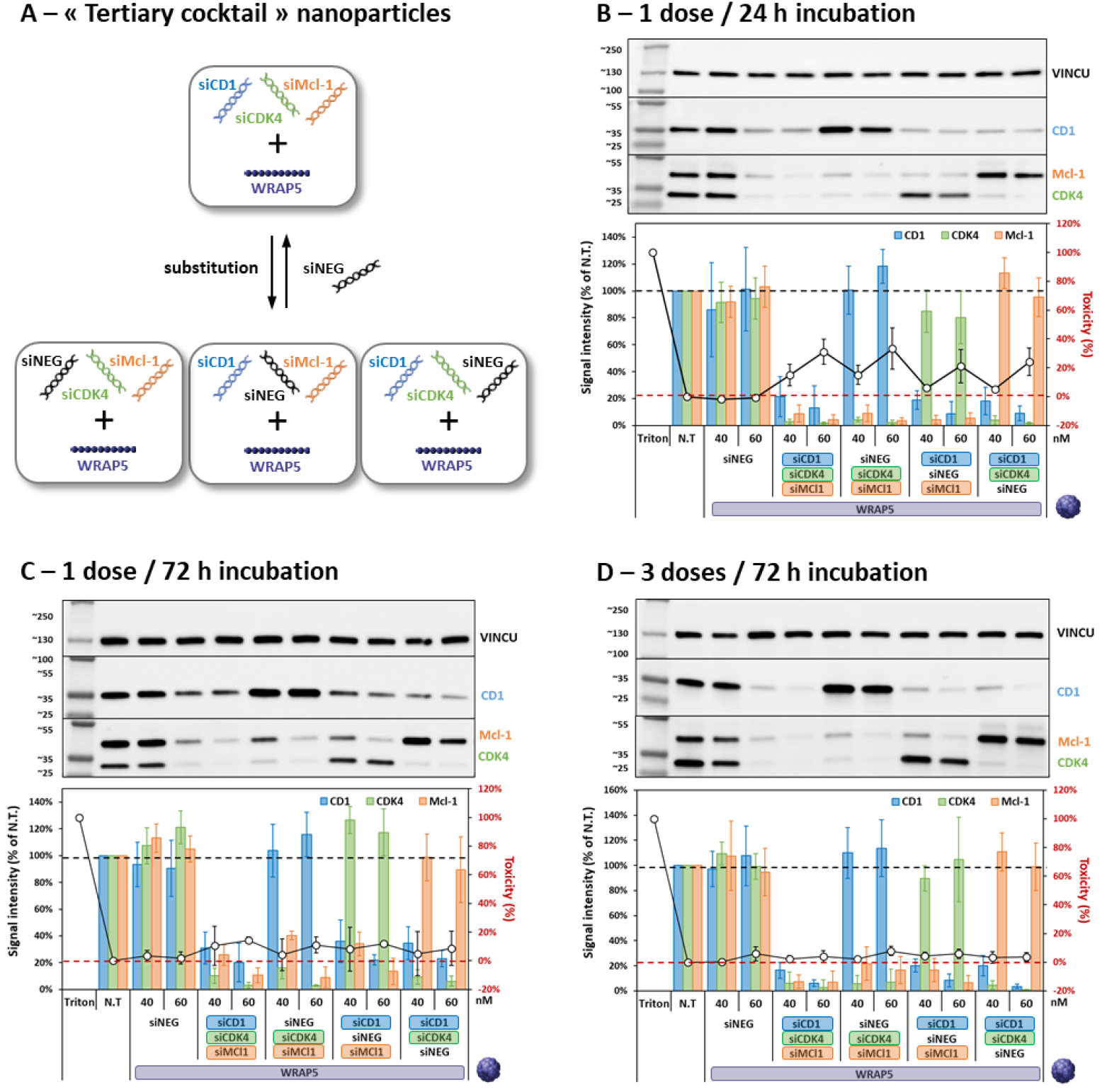
Evaluation of the dose-dependent silencing of WRAP5:[siCD1/siCDK4/siMCL-1] (14/3/3) nanoparticles. **(A)** Schematic representation of WRAP5 nanoparticles encapsulating three siRNAs with or without siNEG substitution. **(B)** Single dose application of WRAP5:[siCD1/siCDK4/siMCL-1] nanoparticles at a ratio 14/3/3 induced a simultaneous silencing of three proteins (CD1, CDK4, and MCL-1) in human U87 glioblastoma cells at the indicated siRNA concentrations revealed by Western Blot after a 24 h incubation. **(C)** Single dose application of WRAP5:[siCD1/siCDK4/siMCL-1] nanoparticles at a ratio 14/3/3 induced a simultaneous silencing of three proteins (CD1, CDK4, and MCL-1) in human U87 glioblastoma cells at the indicated siRNA concentrations revealed by Western Blot after a 72 h incubation. **(D)** Three-dose application of WRAP5:[siCD1/siCDK4/siMCL-1] nanoparticles at a ratio 14/3/3 induced a simultaneous silencing of three proteins (CD1, CDK4, and MCL-1) in human U87 glioblastoma cells at the indicated siRNA concentrations revealed by Western Blot after a 72 h incubation. Data represent mean ± SD, with n = 3 independent experiments in duplicates. Non-treated cells (N.T.) / Triton were used to determine 100% toxicity using LDH assay. The dashed black line represented 100% protein expression and the dashed red line represented 0% cytotoxicity

When applying one single dose of the WRAP5:siCOCK nanoparticles, we observed after 24 h incubation of the U87 cells a clear silencing (≥80%) for the three proteins which was not observed with the WRAP5:siNEG (**Figure 2B**). Moreover, if one siRNA of the cocktail was replaced by the siNEG, the corresponding protein was expressed in the same manner as in the non-treated cells, confirming the specificity of the WRAP5:siCOCK. For the 60 nM siCOCK concentration, we could observe a slight cell cytotoxicity (20%-40%) which was not shown for the WRAP5:siNEG at the same concentration. The results demonstrated that WRAP5:siCOCK nanoparticles appeared to alter cell survival mechanisms, regardless of whether two or three siRNAs were encapsulated.

Next, we assessed the duration of the silencing effect of a single dose of WRAP5:siCOCK by analyzing the protein expression of CD1, CDK4, and MCL-1 at 72 h post-transfection. Indeed, the silencing observed at 24 h was maintained over time, revealing a still important knock-down of more than 70% of all three proteins (**Figure 2C**). However, the previously observed cytotoxicity (>20%) was lost probably due to the low stability of the lactate dehydrogenase (LDH) enzyme over the 72 h of incubation.

We then investigated whether using WRAP5:siCOCK nanoparticles at a reduced concentration, in conjunction with repetitive doses, would have any further impact on protein silencing. To this end, we applied lower concentrations of siCOCK (20 and 40nM) daily for three consecutive days and observed the effects after 72 h post-transfection. As shown in **Figure 2D**, the results indicated that three repeated doses of WRAP5:siCOCK induced the same silencing effect on the three targeted proteins (CD1/CDK4/MCL-1) as observed for the single dose while using two-fold lower concentrations (20 nM and 40 nM) (**Figure 2C**). However, no specific cytotoxic effects were observed due to the siCOCK incubation, even with the three doses applied. At the time of the third treatment (48 h), cells were still proliferating resulting in a higher cell density compared to the conditions at 24 h. It is known that the possible cytotoxicity of nanoparticles depends on the cell density in the well: Cells with a lower density are more affected than cells with a higher density.^33^

### WRAP5:siCOCK nanoparticles: impact on U87 cell proliferation

As we observed a slight increase of cytotoxicity with the WRAP5:siCOCK nanoparticles (**Figure 2A**), we aimed to compare them with the WRAP5 nanoparticles encapsulating the single siRNA. For that, the siRNA concentrations within the “single”-loaded nanoparticles corresponded to those used in the “tertiary siCOCK” using a short (24 h) and a longer (14 days) incubation period.

First, we evaluated the 24 hour condition and revealed efficient silencing (~80%) of the three proteins using the WRAP5:siCOCK nanoparticles (**Figure 3A**), comparable to the knockdown observed above in **Figure 1C**. This important silencing (80%) was only observed with WRAP5 nanoparticles loaded with the single siCD1 with the highest siRNA concentration (28 nM). For the two other single siRNA-loaded WRAP5 nanoparticles (siCDK4 and siMCL-1) the knock-downs reached lower values (60% and 45% for CDK4 and MCL-1, respectively) compared to siCOCK, supporting synergistic effects. In addition, all observed protein silencing phenomena were specific because WRAP5:siNEG gave nearly no knock-down activity (100% ± 20%) for the three targeted proteins. Moreover no significant levels of cellular toxicity was detected by LDH after 24 hours.

**Figure 3:**
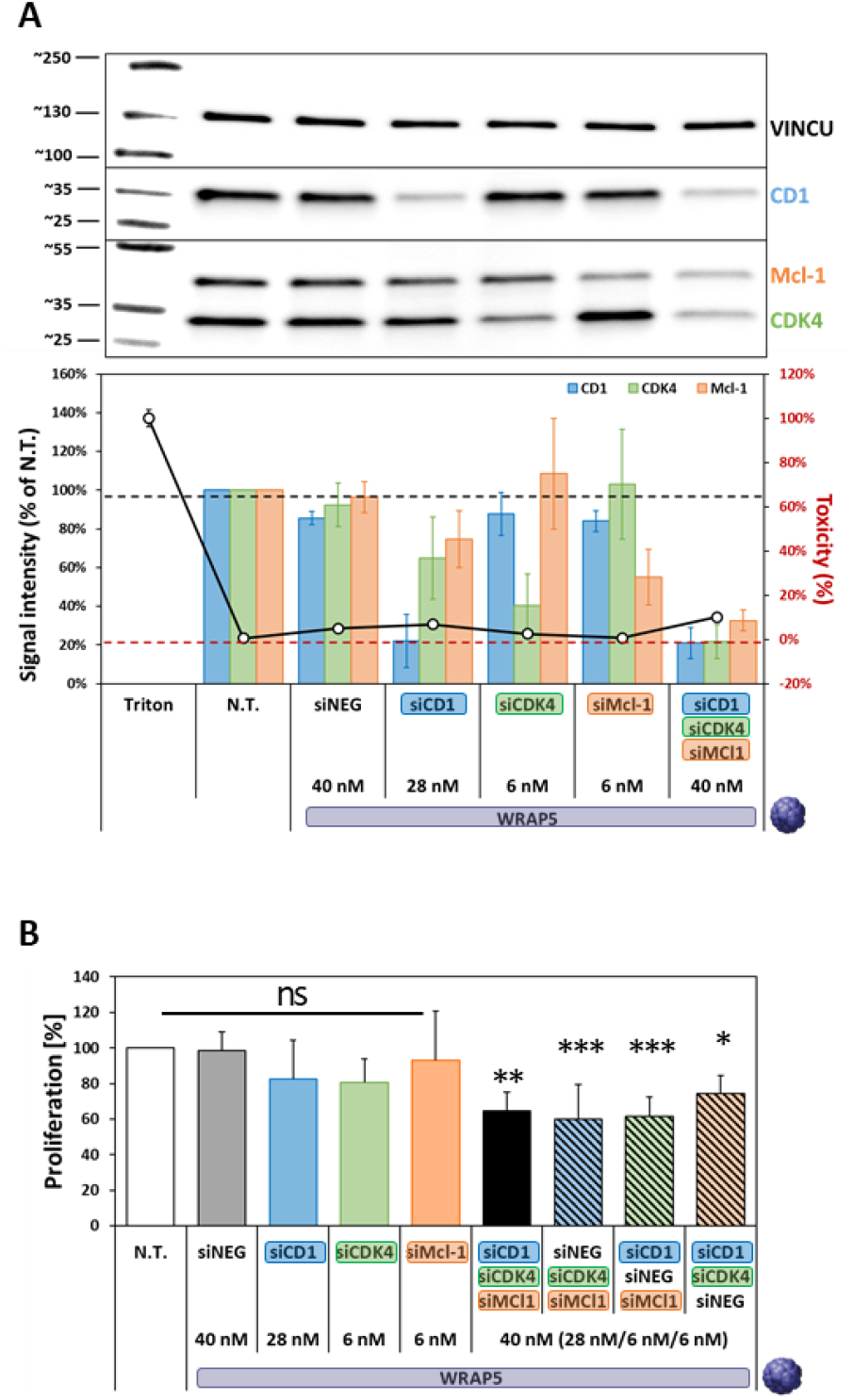
WRAP5:siCOCK (14/3/3) nanoparticles affect cell proliferation on U87 cells depending on the incubation time. **(A)** WRAP5:[siCD1/siCDK4/siMCL-1] nanoparticles at a ratio 14/3/3 induced a simultaneous silencing of three proteins (CD1, CDK4, and MCL-1) in human U87 glioblastoma cells at the indicated siRNA concentrations revealed by Western Blot after a 24 h incubation. Data represent mean ± SD, with n = 2 independent experiments in duplicates. Non-treated cells (N.T.) / Triton were used to determine 100% toxicity using the LDH assay. The dashed black line represented 100% protein expression and the dashed red line represented 0% cytotoxicity. **(B)** WRAP5:[siCD1/siCDK4/siMCL-1] nanoparticles at a ratio 14/3/3 induced a reduction in cell proliferation in human U87 glioblastoma cells at the indicated siRNA concentrations revealed by clonogenic assay after a 14 days incubation. Data represent mean ± SD using >5 independent experiments. Statistics: Kruskal-Wallis test ns >0.05, * p <0.05, ** p <0.01, *** p <0.001, **** p <0.0001 *versus* N.T.

Then, we compared the effect of WRAP5:siCOCK nanoparticles on a long (14 days) incubation period using a clonogenic assay (**Figure 3B**). The clonogenic assay is a well-established experimental approach that quantifies the ability of a single cell to form a colony. It is a widely utilized technique for assessing the effects of pharmaceutical agents on the growth and proliferative characteristics of cells *in vitro*.

First, we showed that the single siRNA-loaded WRAP5 nanoparticles (same concentration as in the siCOCK) had no significant effect on cell proliferation compared to the non-treated or the WRAP5:siNEG-treated cells. More interestingly, WRAP5:siCOCK (40 nM) nanoparticles demonstrated a significant proliferation reduction of ~40% (** p <0.01). Afterward, we evaluated the potent synergic effect of the three siRNAs within the siCOCK by replacing each of them with the siNEG as shown previously (**Figure 2B**). When siCD1 or siCDK4 was replaced by the siNEG, we observed a comparable reduction in cell proliferation as the WRAP5:siCOCK condition. However, in the case of replacing the siMCL-1 by the siNEG, cell proliferation was less reduced (20%, * p <0.05), suggesting that, compared to siCDK4 and siCD1, the “tertiary cocktail” required siMcl-1 to ensure a greater impact. We hypothesized that the resulting inhibition of Mcl-1 could be correlated to apoptosis induction, as the knockdown of *Mcl-1* was already associated with an increased level of apoptotic markers.^34^

In conclusion, applying a long-term incubation of WRAP5:siCOCK nanoparticles to the U87 cells we could observe for the first time a physiological effect of the WRAP5-mediated silencing of three proteins resulting in the significant reduction of proliferation.

### WRAP5:siCOCK nanoparticles: suitable for GIST cell transfection

In order to evaluate the universal potential of the “tertiary cocktail” approach, we considered the multiple protein targeting using WRAP-based nanoparticles in another cancer type, namely the gastrointestinal stromal tumor (GIST). To this aim, we first examined the ability of WRAP5 nanoparticles to deliver each siRNA targeting CDK4, CD1, and MCL-1 (siCD1, siCDK4, and siMCL-1) individually in human GIST-T1 cells.^35,36^ Although significant knock-downs were detected for siCD1 and siMCL-1 (>90% for 60 nM), the targeting of CDK4 seemed to reach a plateau of 60% silencing whatever the applied siCDK4 concentration (40, 60, 80 nM), suggesting a different silencing behavior of CDK4 (**Figure S3**). The siCDK4 has been previously validated in human U87 glioblastoma cells ^16,31,32^ but this is the first instance of its use in GIST-T1. Consequently, we tried additional CDK4-targeting siRNAs to identify the most effective siCDK4 for GIST-T1 cells. Three new siRNAs targeting CDK4 were compared to the first, which finally appeared to be the most suitable (**Figure S4**).

Having the optimal siRNA to silence CD1, CDK4, and MCL-1 in human GIST-T1 cells, our next objective was to examine the capacity of WRAP5 nanoparticles to deliver the siCOCK in human GIST-T1 cells using an equivalent ratio (1/1/1) to determine if the three siRNAs were internalized in the same way. For this purpose, WRAP5:siCOCK was formulated using fluorescence-labeled siRNA (siCDK4-Alexa488, siCD1-Atto550 and siMCL-1-Atto-655). Nanoparticle formulations in the presence of fluorescence-labeled siRNA were checked by DLS measurements in terms of size and stability and compared to non-florescence-labeled siRNA (**Table 1**). All nanoparticles formulated with fluorescence-labelled siRNA revealed a size between 75-100 nm with a PdI <0.3 which was more or less equivalent to the unlabeled nanoparticles. We observed a slight discrepancy for the nanoparticle WRAP5:siCD1-Atto550 probably due to the fluorescent properties of the Atto550 compared to green laser features of the Zetasizer NanoZS (Malvern). Indeed the green laser of the DLS produced a 532 nm excitation (50 mW) which could slightly excite Atto550 whose absorbance spectrum starts at 520 nm with a maximum/optimal excitation at 550 nm. This slight excitation could result in a signal noise increase and a higher polydispersity index (***Malvern FAQ website***).^37^

The internalization of WRAP5:siCOCK nanoparticles encapsulating the three fluorescence-labeled siRNAs was visualized by confocal microscopy, thereby revealing a visible cytoplasmic signal in nearly all transfected cells compared to the non-treated cells or those treated with the siCOCK without WRAP5 as the delivery system (**Figure 4A**). Live-cell imaging revealed the typical dotter pattern of internalized siRNA (siCD1-Atto550, siCDK4-Alexa488, and siMCL-1-Atto654) visible in the cytoplasm of GIST-T1 cells as previously reported for human U87 glioblastoma cells.^16,17^

**Figure 4:**
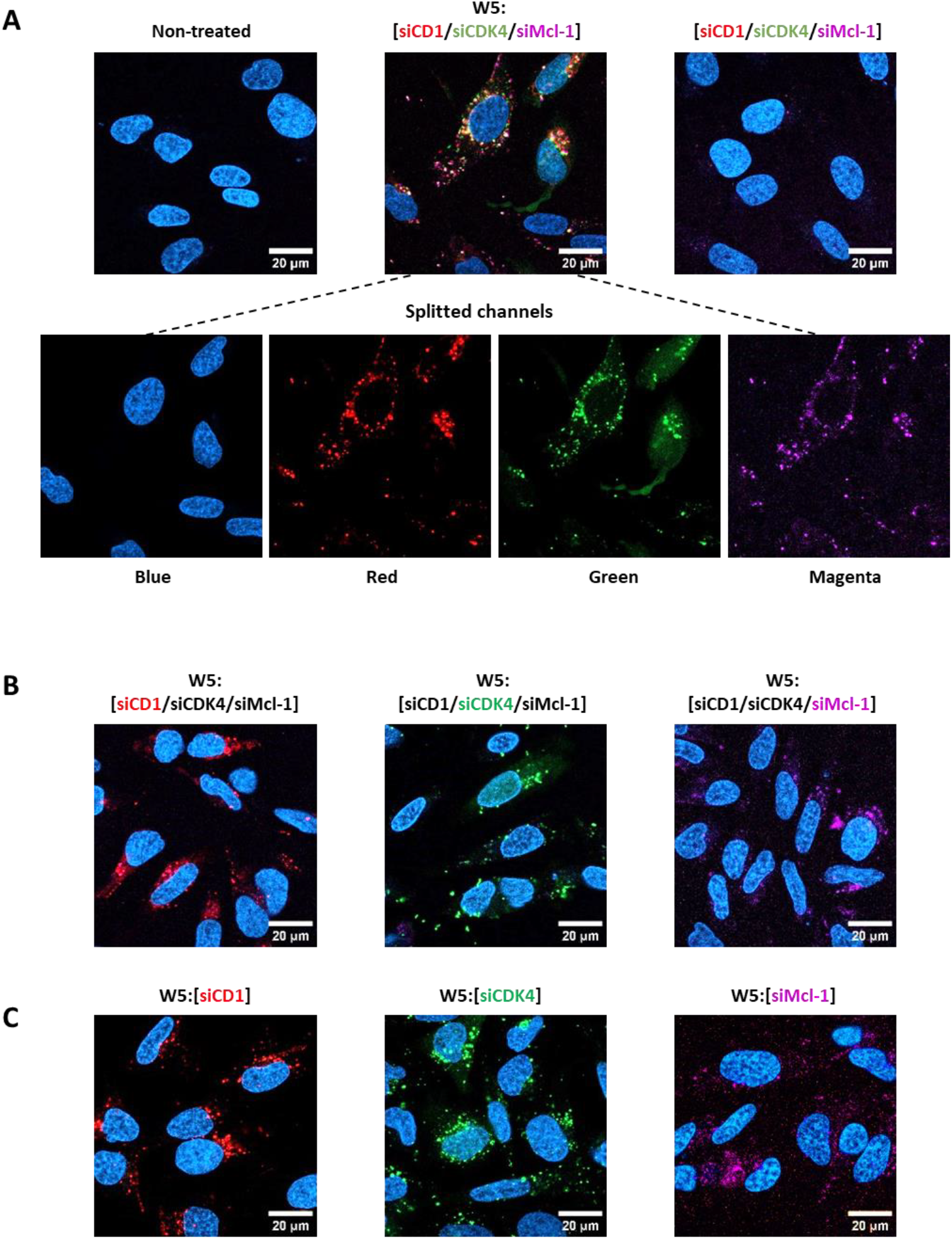
Evaluation of WRAP5:[siCD1/siCDK4/siMCL-1] nanoparticles transfection in GIST-T1 cells. (**A**) Representative live images of confocal microscopy acquisitions of fully fluorescence-labeled WRAP5:[siCD1/siCDK4/siMCL-1] nanoparticles (ratio 1/1/1, with 40 nM total siRNA concentration) compared to non-treated GIST cells or those treated with the siRNA alone [siCD1/siCDK4/siMCL-1] (without WRAP5). As a control for the uniform cellular internalization, representative live images of confocal microscopy acquisitions WRAP5:[siCD1/siCDK4/siMCL-1] nanoparticles (ratio 1/1/1, with 40 nM total siRNA concentration) with only one fluorescence-labeled siRNA (**B**) compared to those incubated the single siRNA-loaded WRAP5 nanoparticles (40 nM) (**C**). Conditions: GIST-T1 cells were incubated for 6 h at the indicated siRNA compositions. Red = siCD1-Atto550, green = siCDK4-Alexa488, magenta = siMCL-1-Atto654 and blue = Hoechst dye. White bar = 20 μm.

Afterward, the internalization of WRAP5:siCOCK nanoparticles containing solely one fluorescence-labeled siRNA out of the three available siRNA was compared to the internalization of WRAP5:siRNA encapsulating siCD1 or siCDK4 or siMcl-1 (**Figure 4B)**. This comparison revealed that the internalization of the labeled siRNAs occurred in a similar manner, irrespective of whether they were encapsulated alone or in combination with the others.

In conclusion, these microscopy results indicated that the presence of the three siRNAs within a single nanoparticle induced simultaneous cell transfection. In all cases, the observed fluorescence signals were specific for WRAP5-based transfection as no fluorescence signal was observed when siCOCK was applied to the cells alone.

### WRAP5:siCOCK nanoparticles: suitable simultaneous protein in GIST cells

Before evaluating the silencing abilities of the WRAP:siCOCK to silence the expression of CDK4, CD1 and MCL-1 in human GIST-T1 cells, we first investigated their basal expression levels in GIST-T1 cells compared to U87 glioblastoma cells. At equal applied total protein concentration, we determined by Western blot analysis an equal expression level for CD1 and MCL-1 proteins but a higher expression pattern for CDK4 in GIST-T1 cells (**Figure 5A**). Due to this fact, we considered that the siRNA ratio in the cocktail should be modified to fit with protein expression profiles in human GIST-T1 cells. We used a higher amount of siCDK4 compared to CD1 and MCL-1 to reach an equal silencing of all three proteins. First, we evaluated the two siRNA ratios of 3/14/3 and 4/12/4 for siCD1, siCDK4, and siMCL-1, respectively (**Figure 5B**), revealing a dose-dependent silencing of the three targets. However, the knockdown of CD1 seemed to be less pronounced than for the other two proteins (CDK4 and MCL-1), suggesting that higher siCD1 concentrations were needed. To consider a higher amount of siCD1 in the cocktail, two other 5/10/5 and 6/8/6 siRNA ratios were evaluated in a dose-dependent manner in GIST-T1 cells (**Figure 5C**). In this context, compared to non-treated or GIST-T1 cells treated with the nanoparticles loaded with siNEG, only the WRAP5 nanoparticles encapsulating the 6/8/6 siRNA ratio revealed a homogeneous silencing of the three proteins of around 50% and 70% at a concentration of 40 nM and 60 nM, respectively. Under these conditions, slight cytotoxicity (LDH assay) was shown at high siRNA concentration independent of the used siRNA (also siNEG) suggesting that this phenomenon was due to the transfection conditions rather than silencing effects.

**Figure 5:**
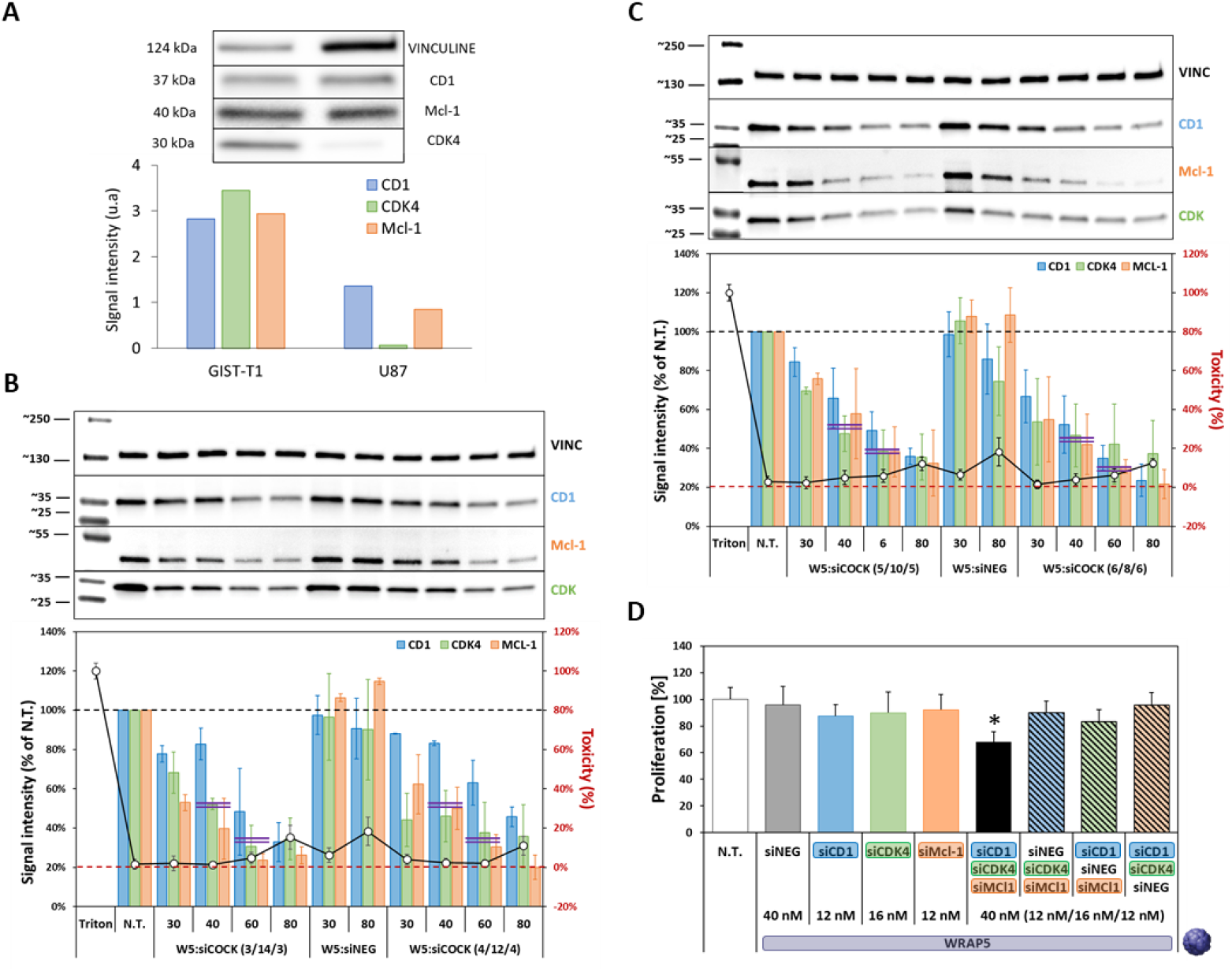
Evaluation of WRAP5:[siCD1/siCDK4/siMCL-1] nanoparticles transfection in GIST-T1 cells. **(A)** Basal expression levels of CD1, CDK4, and MCL-1 proteins in human GIST-T1 compared to U87 cells. We determined an equal expression level for CD1 and MCL-1 proteins but a higher expression pattern for CDK4 in GIST-T1 cells, contrary to U87 cells, where CDK4 was less expressed than CD1 and MCL-1. **(B)** Comparison of single-dose application of WRAP5:[siCD1/siCDK4/siMCL-1] nanoparticles at a ratio 3/14/3 or 4/12/4 at the indicated siRNA concentrations revealed by Western Blot after a 24 h incubation. Both conditions induced a simultaneous silencing of CD1 and MCL-1 proteins in GIST cells with less efficiency for CD1 protein (see magenta double bars). Data represent mean ± SD using 3 independent experiments. **(C)** Comparison of single-dose application of WRAP5:[siCD1/siCDK4/siMCL-1] nanoparticles at a ratio 5/10/5 or 6/8/6 at the indicated siRNA concentrations revealed by Western Blot after a 24 h incubation. Only the condition with a siRNA ratio of 6/8/6 induced a simultaneous silencing of all three proteins in GIST cells (see magenta double bars). Data represent mean ± SD using 3 independent experiments. (D) WRAP5:[siCD1/siCDK4/siMCL-1] nanoparticles at a ratio 6/8/6 induced a reduction in cell proliferation in U87 cells at the indicated siRNA concentrations revealed by clonogenic assay after a 14 days incubation. Data represent mean ± SD using >3 independent experiments. Statistics: Kruskal-Wallis test * p <0.05 versus non-treated cells (N.T.).

### WRAP5:siCOCK nanoparticles: suitable to reduce cell proliferation in GIST-T1 cells

Finally, as described for U87 cells, the long-term impact of the WRAP5:siCOCK nanoparticles on cell proliferation was investigated in human GIST-T1 cells through a similar clonogenic assay with a 14 days incubation period (**Figure 5D**). First, we analyzed the WRAP nanoparticles encapsulating the single siRNA, showing no effect on GIST-T1 cell proliferation. Then, we selected WRAP5:siCOCK nanoparticles at the 6/8/6 siRNA ratio for siCD1, siCDK4, and siMCL-1, respectively, as the optimal transfection condition showing the highest knock-down efficiency. Using this condition, we demonstrated a significant reduction of cell proliferation of ~30% (* p <0.05). Interestingly, this WRAP:siCOCK appeared as the only condition showing a physiological impact on GIST-T1 cells. Indeed, nanoparticles encapsulating siNEG substitution did not induce any effect on GIST-T1 colony formations (**Figure 5D**).

In conclusion, these results revealed that a physiological effect could be only detected for the simultaneous silencing of three proteins in GIST-T1 cells through a long-term incubation (14 days without medium change) of WRAP5:siCOCK nanoparticles. Furthermore, the difference in silencing of the three targets was found to be associated with the discrepancy of physiological impact (cell proliferation) between human U87 and GIST-1-T1 cells. This emphasized the importance of adapting siRNA ratio in siCOCK according to the cell types and the expression profile of the corresponding targets.

## DISCUSSION

### WRAP5-based siRNA delivery in human U87 glioblastoma cells: from one to multiple silencing

First, we investigated the use of WRAP5-based nanoparticles to transfer siRNA such as siCDK4, siCD1, and siMCL-1 individually in human U87 glioblastoma cells to estimate the efficiency of gene silencing for each target and their potential physiological impact (**Figure 1A**). As already published, we observed that the targeting of CDK4 was significant (95% inhibition with 20 nM siCDK4) and in agreement with our previous work ^16^, supporting here the robustness of the WRAP5 technology in siRNA delivery. The siRNA-mediated inhibition of CD1 expression was significant but required a higher siRNA concentration (50 nM) to reach a similar CDK4 extinction, suggesting a difference in the level of expression or half-life of CD1 compared to CDK4. However, as for CDK4, the CD1 inhibition did not induce any significant phenotype. Concerning MCL-1, a strong knockdown (85% inhibition) was detected at 20 nM. Belonging to the anti-apoptotic BCL-2 family proteins, MCL-1 was known to have a very short biological half-life of only 20-30 minutes, which could explain the siRNA-mediated knock-down efficiency.^38^ However, we did not detect any impact on the cell proliferation for the three single siRNAs used in this study. We correlated this observation with the fact that CDK4, as well as CD1, could be replaced by other CDK or Cyclin proteins.^27,39^

In the context of glioblastoma cells, it is known that Bcl-2 inhibition triggers MCL-1 upregulation. This compensatory mechanism is a cellular response to the role typically played by Bcl-2 in preventing cell death.^40^ Thus, it is possible that MCL-1 anti-apoptotic activity could be replaced by other anti-apoptotic proteins such as Bcl2.

As for other drug delivery systems, we tried to improve the cellular impact of gene silencing by using siRNA combinations through WRAP5 nanoparticles encapsulating siRNA cocktails. We first investigated a “binary cocktail” approach by encapsulating two siRNAs within the nanoparticle. We tested the association of siCD1 with siCDK4 to knockdown the CDK4/CD1 complex involved in the cell cycle checkpoint of human U87 glioblastoma cells, expecting a cell cycle arrest. As the CD1 inhibition alone required a higher amount of siRNA, we mixed 80% of siCD1 with 20% of siCDK4, corresponding to a 4/1 stoichiometry between siCD1/siCDK4 (**Figure 1B**). Results revealed the ability to silence both proteins at a similar level for this siRNA molar ratio, confirming the rationale of the “binary cocktail” strategy. However, as with previous single-target extinction, no physiological impact was observed, suggesting that potential cell cycle arrest may not be sufficient to generate a net biological effect.^39^

Then, we decided to incorporate a siRNA targeting the anti-apoptotic MCL-1, to orient the approach towards a “tertiary cocktail” strategy with two siRNAs targeting the cell cycle and one targeting anti-apoptotic characteristics (**Figure 1C**). The idea was to affect the cell cycle progression and simultaneously activate apoptosis. Based on a single siRNA and “binary cocktail” formulation, we used a siRNA cocktail with a higher siCD1 concentration (70%) compared to equivalent amounts of both siCDK4 (15%) and siMCL-1 (15%), resulting in a 14/3/3 stoichiometry (70%/15%/15%). The WRAP5-mediated delivery of such a “tertiary cocktail” enabled an equivalent knock-down of the three targets CD1, CDK4, and MCL-1, revealing the force of the WRAP5 nanoparticles as a delivery system.

Using the WRAP5:siCOCK, we observed an 80% extinction of the three respective protein targets at the low siRNA concentration (40 nM siCOCK = 28 nM siCD1, 6 nM siCDK4, 6nM siMCL-1), compared to the 40% to 60% extinction detected for the siRNA individually. These data indicated a synergic effect between the three siRNAs of the siCOCK, supporting the potency of the “tertiary cocktail” combination for multiple silencing by WRAP5 nanoparticles. Our results were in agreement with previously reported nanoparticles loaded with different siRNAs. One example is given through a cholesterol-modified antimicrobial peptide (AMP) DP7, which encapsulated multiple siRNA elements and exhibited a synergic antitumor effect compared to single-target therapy.^41^ Also, dendrimers were shown to be able to encapsulate three distinct siRNAs targeting the *Bcl-xl, Bcl-2*, and *MCL-1* genes for simultaneous protein silencing as a new treatment for breast cancer.^42^ Similarly, lipid nanoparticles were associated with siRNA cocktails enabling inhibition of *MCL-1, Bcl-2*, and *CyclinD1* in the Mantle cell lymphoma.^39^

Finally, we evaluated the impact of WRAP5:siCOCK on the proliferation of human U87 glioblastoma cells using the commonly used clonogenic assay. We first demonstrated that single siRNA-loaded nanoparticles were ineffective in altering cell proliferation. Remarkably, we highlighted that WRAP5:siCOCK nanoparticles significantly reduced cell proliferation (−40% with 40 nM) underscoring the importance of multi-targeting genes for effective cancer inhibition.

### siRNA cocktail optimization based on targeted cancer cell types

After validation in human U87 glioblastoma cells, we applied our multiple targeting strategies to other cancers by considering gastrointestinal stromal tumors (GIST), a common form of sarcoma of the digestive tract. A rapid comparison of the basal expression levels of targeted proteins revealed a difference in the CDK4 expression profile with a higher level in GIST-T1 than in U87 cells. To translate the WRAP5:siCOCK to human GIST-T1 cells modification of the siRNA molar ratio within the siCOCK should be changed in order to reach a similar silencing efficiency.

This first observation was not surprising as cancer cell types have different properties and protein expression suggesting that the siRNA ratio should be adapted. An example is given by Knapp et al. mentioning the treatment of human mantle cell lymphoma (JeKo-1 cells) using a “tertiary siCOCK” at a dose of 210 nM, which contained 10 nM siMCL-1, 100 nM siBcl-2, and 100 nM siCD1 (1/10/10 siRNA molar ratio).^39^ In our study, a low siMCL-1 concentration was sufficient in comparison to the siCD1/siBcl-2 concentrations.

For GIST-T1 cells, we used the same three siRNAs (siCD1, siCDK4, and siMCL-1) as all were represented in this cell line associated with the development of the pathology. CD1 was described as a potential therapeutical target in KIT-independent GISTs, with bortezomib-mediated inhibition proposed as a novel therapeutic strategy.^43^ Furthermore, CD1 played a crucial role in GIST-KIT independence, as its suppression exhibited anti-proliferative and pro-apoptotic effects.^30^ Besides that, recurrent genomic aberrations by in-cell cycle regulators were determined which caused co-activation of the CDK2 and CDK4/6 pathways in clinical GIST samples.^27^ Finally, MCL-1 overexpression in GIST patients was associated with OSTEOPONTIN, a secreted phosphoprotein involved in the malignant potential and aggressive phenotypes.^44^

After having screened several siRNA ratios, we identified the ratio of 6/8/6 [siCD1/siCDK4/siMCL-1] as the optimal one for human GIST-T1 cells (**Figure 5C**). In detail, we observed with this optimized siCOCK the silencing of around 50% (40 nM) of the three proteins CD1, CDK4, and MCL-1. Curiously, using the same siCOCK concentration of 40 nM, the knockdown efficiency in GIST-T1 cells was lower than in human U87 glioblastoma cells. Clonogenic assay on GIST-T1 cells using WRAP5:siCOCK nanoparticles revealed a significant reduction (−30% with 40 nM) compared to single siRNA or siNEG-substituted siCOCK highlighting their impact on GIST-T1 cell proliferation (**Figure 5D**).

These results supported the fact that a stoichiometric siRNA adjustment should be performed according to the cellular type, i.e. targeted cancer. This also indicated that for similar targeted proteins, the innate nature of cells determined the siRNA cocktail stoichiometry.

## CONCLUSION

In the present work, we demonstrated the ability of WRAP5-based nanoparticles to deliver a siRNA cocktail (siCOCK) in order to silence three distinct proteins in neuroblastoma and sarcoma-derived cells. The siRNA molar ratio within the siCOCK had to be adapted to the level of expression of each targeted protein with regard to the cancer cell type used. However, working on siRNA cocktails encapsulated in delivery systems raised several questions as highlighted by Guo F. et al.^45^ Before the selection of the targeted proteins that should be silenced, researchers should ask themselves if multigene silencing will be more efficient than single gene knockdown, if targeting different signaling pathways should be more suitable than focusing on a single pathway. Furthermore, it is important to identify the main target/siRNA couple to define the optimal combination (ratio) that should be applied to obtain the best synergic therapeutic gene silencing.

For the future, to potentiate WRAP5-based siRNA cocktails, other parameters could be further optimized as reported previously to increase their *in vivo* stability by PEGylation ^32^, their localization in specific intracellular compartments by targeting motif grafting ^46^ or more recently their dependence on the nature of the microenvironment by integration of a pH-sensitive linker.^47^

Taking together, these innovative results with respect to peptide-based nanoparticle technology will pave the way for developments that could lead to an improvement in the approach to personalized medicine.

## ASSOCIATED CONTENT

### Supporting Information

The authors have included the whole Experimental Section, Tables S1, and Figures S1 to S4 in the Supporting Information. Table S1 referred to the used siRNA sequences; Figure S1 to the evaluation of the dose-dependent silencing of WRAP5:siRNA nanoparticles in U87 cells; Figure S2 to graphical representations of the measured mean sizes of WRAP5-based nanoparticles; Figure S3 to the evaluation of the dose-dependent silencing of WRAP5:siRNA nanoparticles in GIST cells; Figure S4 to the evaluation of the silencing of WRAP5:siRNA nanoparticles encapsulating different siCDK4 in GIST-T1 cells

## ACKNOWLEDGMENTS

This research was funded by the “Fondation ARC pour la Recherche sur le Cancer” (PJA20171206171 to P.B.), the “Agence Nationale de la Recherche” (ANR-21-CE18-0022-01 to P.B.), AFM-Téléthon (n°23800 to S.F.), and by institutional funds from INSERM, CNRS and University of Montpellier. The authors are grateful to Pascal Verdié from the SynBio3 platform for providing peptide synthesis facilities, and to Pierre Sanchez for performing LCMS analysis, both from the Institut des Biomolécules Max Mousseron (IBMM), Montpellier (France).

## Declaration of Interests

Patent entitled “Non-naturally occurring peptides for use as cell-penetrating peptides” WO2020/016242.

## Conflicts of Interest

The authors declare no competing interests.

## Notes

### Competing Interest Statement

The authors have declared no competing interest.

