## Supplement Material for "Multi-protein silencing using WRAP-based nanoparticles: a proof of concept"

**SUPPORTING INFORMATION**

**Table S1: Used siRNA sequences**

| ID | Anti-sense sequences | References |
|---|---|---|
| **siCDK4*** | 5′-CAG-AUC-UCG-GUG-AAC-GAU-GdTdT-3′ | Mao 2014 [1] |
| **siCDK4 (2)** | 5′-UGU-GGG-UUA-AAA-GUC-AGC-AdTdT-3′ | Ikai 2016 [2] |
| **siCDK4 (3)** | 5′-CUC-CAU-CUU-UCU-ACA-GAU-UAC-dTdT-3′ | Kathryn 2013 [3] |
| **siCDK4 (4)** | 5′-CUC-UUA-UCU-ACA-UAA-GGA-UGA-AGG-dTdT-3′ | Kathryn 2013 [3] |
| **siCD1** | 5′-AAA-UGA-ACU-UCA-CAU-CUG-UdTdT-3′ | Champagne 2020 [4] |
| **siMCL-1** | 5′-GGA-CUU-UUA-UAC-CUG-UUA-UdTdT-3′ | Dzmitruk 2015 [5] |
| **siSCR** | 5′-CAU-CAU-CCCUGC-CUC-UAC-UdTdT-3′ | Konate 2019 [6] |

Footnotes: * corresponds to siCDK4 (1) in Figure S4 and was used in all cocktails formulations

WRAP5 nanoparticles encapsulating two siRNA : binary « cocktail »

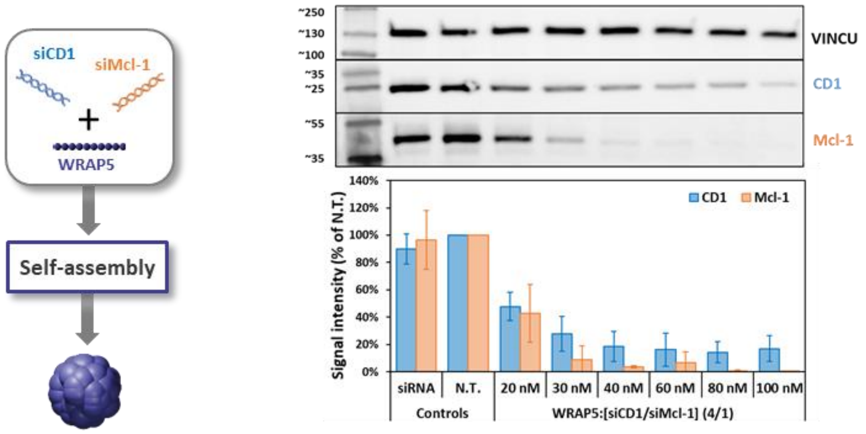


**Figure S1: Evaluation of the dose-dependent silencing of WRAP5:siRNA nanoparticles in U87 cells**
WRAP5 nanoparticles encapsulating two siRNA (schematic representation on the left side) in a final siRNA ratio 4/1 induced the dose-dependent simultaneous silencing of two proteins (CD1/MCL-1) in U87 cells at the indicated siRNA concentrations revealed by Western Blot after a 24 h incubation. Graphical representation of mean ± SD using 3 independent experiments.

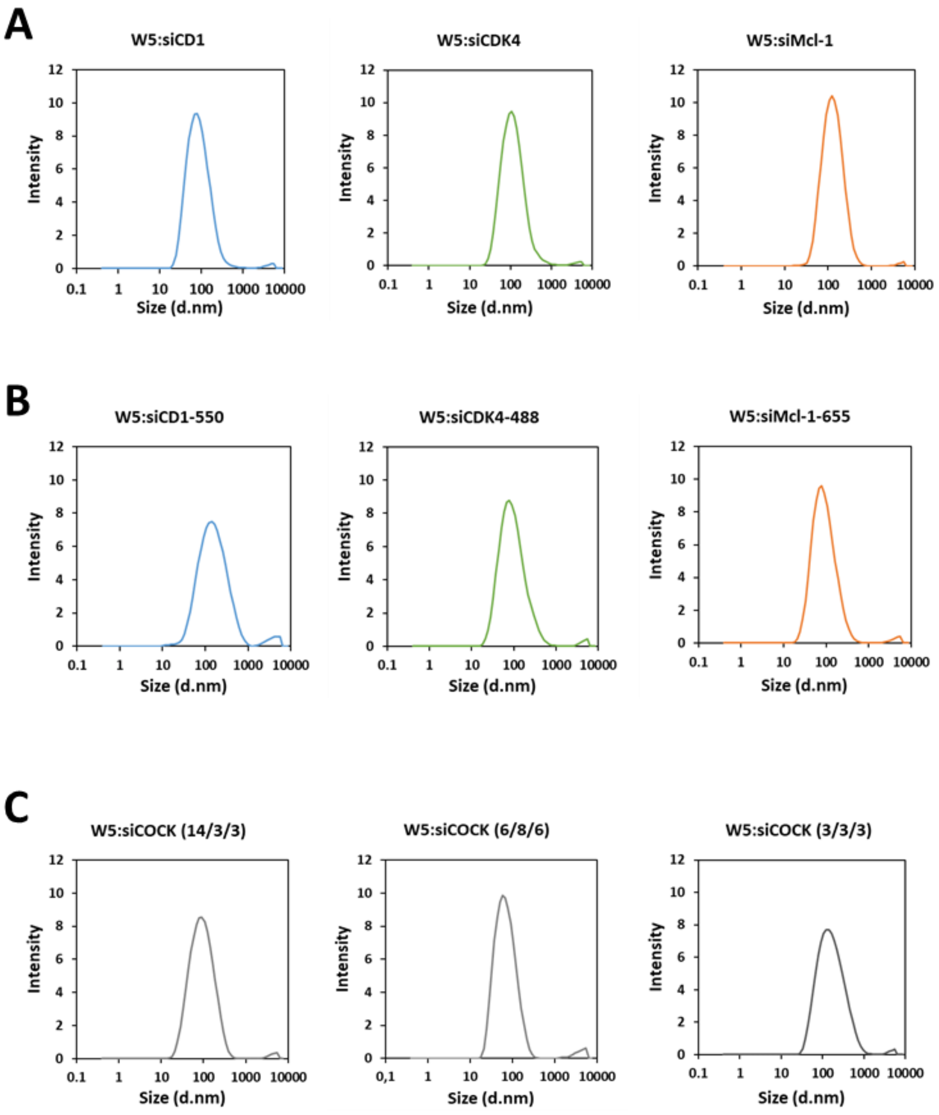


**Figure S2: Graphical representations of the measured mean sizes of WRAP5:siRNA nanoparticles**
Mean size distribution of WRAP5:siRNA nanoparticles (W5:siRNA) for **(A)** single siRNA (siCD1, CDK4, MCL-1) **(B)** single fluorescence-labeled siRNA (siCD1-550, siCDK4-488, siMCL-1-655) and **(C)** siCOCK with siRNA ratio at 14/3/3, 6/8/6, and 3/3/3 molar ratio.

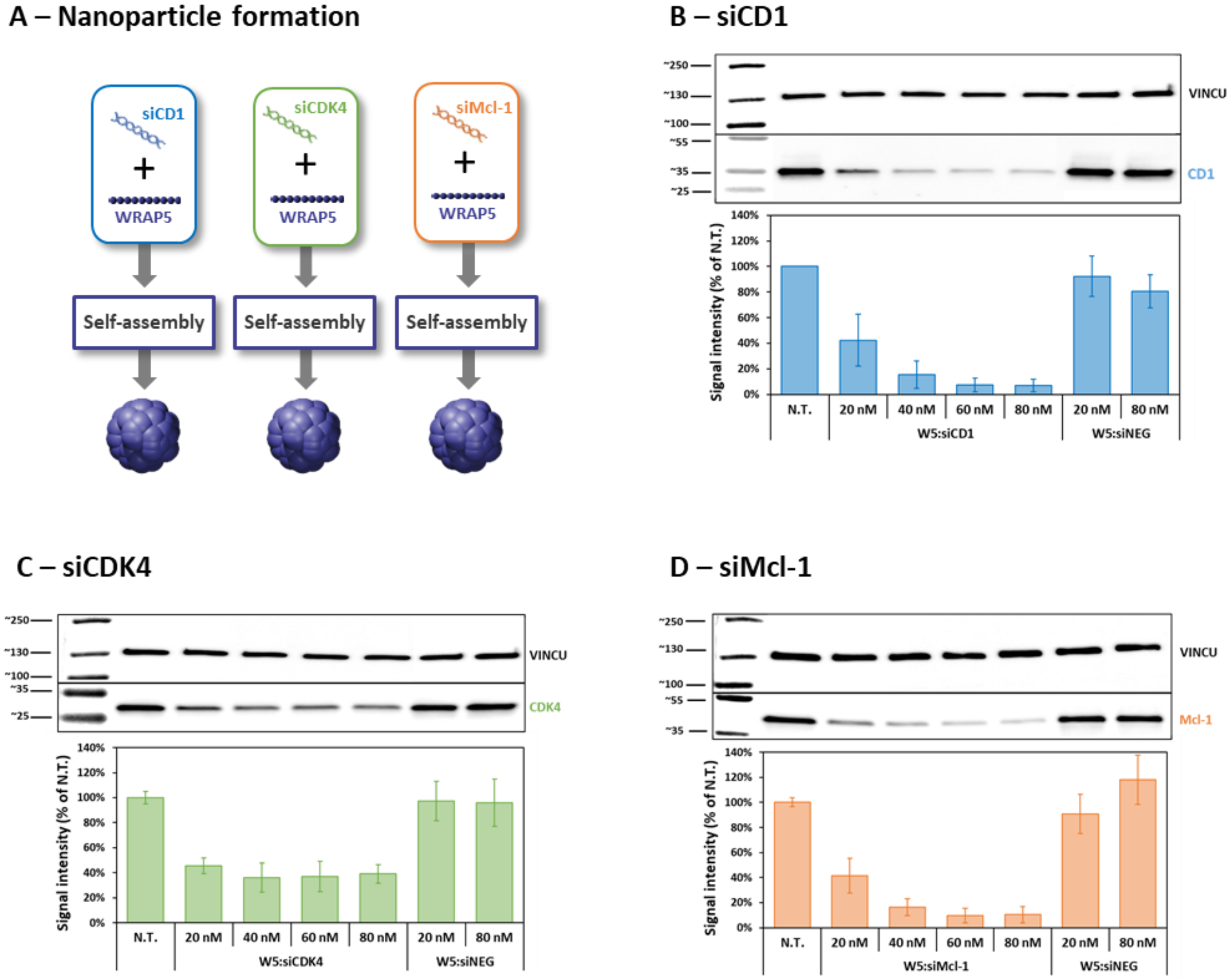


**Figure S3: Evaluation of the dose-dependent silencing of WRAP5:siRNA nanoparticles in GIST-T1 cells**

(**A**) Schematic representation of the nanoparticle formulation using WRAP encapsulating one siRNA. siCD1 (**B**), siCDK4 (**C**) and siMCL-1 (**D**).
Graphical representation of mean ± SD using 3 independent experiments.

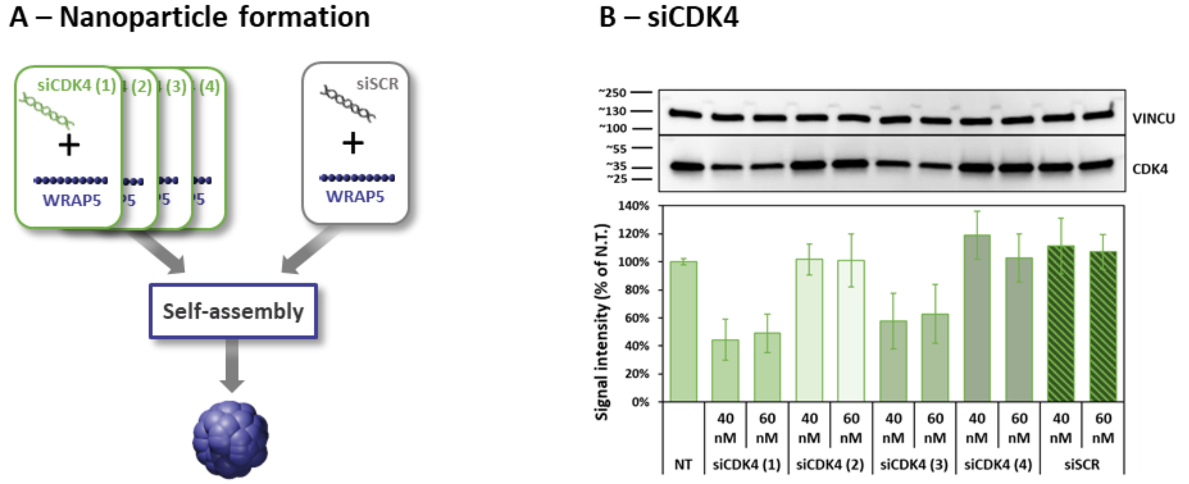


**Figure S4: Evaluation of the silencing of WRAP5:siRNA nanoparticles encapsulating different siCDK4 in GIST-T1 cells**

(**A**) Schematic representation of the nanoparticle formulation using WRAP encapsulating one siRNA.

(**B**) Four different siRNA targeting CDK4 (siCDK4(1), siCDK4(2), siCDK4(3) and siCDK4(4)) were encapsulated in WRAP5-based nanoparticles at both 40 nM / 60nM siRNA concentration in order to check their targeting efficiency in GIST-T1.

Graphical representation of mean ± SD using 2 independent experiments.
